# *Wolbachia* strain wMelM disrupts egg retention by *Aedes aegypti* females prevented from ovipositing

**DOI:** 10.1101/2024.06.12.598754

**Authors:** Perran A. Ross, Ella Yeatman, Xinyue Gu, Ary A. Hoffmann, Belinda van Heerwaarden

**Author notes:** co-senior authors (Ary A. Hoffmann, Belinda van Heerwaarden).

## Abstract

*Aedes aegypti* mosquitoes are well adapted to dry climates and can retain their eggs for extended periods in the absence of suitable habitat. *Wolbachia* strains transferred from other insects to mosquitoes can be released to combat dengue transmission by blocking virus replication and spreading through populations, but host fitness costs imposed by *Wolbachia*, particularly under some environments, can impede spread. We therefore assessed the impact of two *Wolbachia* strains being released for dengue control (*w*AlbB and *w*MelM) on fecundity and egg viability following extended egg retention (12 or 18 d) under laboratory conditions. Egg viability decreased to a greater extent in females carrying *w*MelM compared to uninfected or *w*AlbB females. Fertility fully recovered in uninfected females following a second blood meal after laying retained eggs, while *w*MelM females experienced only partial recovery. Effects of *w*MelM on egg retention were similar regardless of whether females were crossed to uninfected or *w*MelM males, suggesting that fitness costs were triggered by *Wolbachia* presence in females. The fecundity and hatch proportions of eggs of *w*MelM females declined with age, regardless of whether females used stored sperm or were recently inseminated. Costs of some *Wolbachia* strains during egg retention may affect the invasion and persistence of *Wolbachia* in release sites where larval habitats are scarce and/or intermittent.

## Introduction

*Aedes aegypti* are widespread in tropical regions with high rainfall but also possess several adaptations which allow them to tolerate dry climates. Their eggs are desiccation tolerant and can remain viable in a quiescent state for several months (Sota and Mogi, 1992). Moreover, at least under laboratory conditions females can retain viable eggs in their bodies when oviposition sites are unavailable, eggs of *Ae. aegypti* and those of at least several other mosquito species can be retained for weeks at a time with little decline in viability (Johnson and Fonseca, 2014, Xue et al., 2005, Judson, 1968). This includes domestic forms of *Aedes aegypti* live in close proximity to humans that have adapted to lay eggs in human-made containers (Kolimenakis et al., 2021). These females typically mate once and store sperm in spermathecae to produce offspring for their entire life (Degner and Harrington, 2016b). Mating is decoupled from blood feeding (League et al., 2021) and gravid females can retain eggs in their ovaries until mating and finding a suitable oviposition site (Bentley and Day, 1989). This provides *Ae. aegypti* with the flexibility to pause their reproduction for extended periods. This ability to retain eggs can be affected by genetic factors (Venkataraman et al., 2023), but it is unclear whether this variation is driven by ecological differences in conditions experienced by populations. Egg retention could also be affected by other inherited factors, notably maternally inherited endosymbionts including *Wolbachia* residing naturally or increasingly being introduced deliberately into mosquitoes for disease control (Ant et al., 2023). Egg dormancy effects have previously been documented for natural *Wolbachia* in *Drosophila melanogaster* (Kriesner et al., 2016).

Any effects of *Wolbachia* endosymbionts deliberately introduced to control arbovirus transmission could influence the success of programs aimed at using endosymbionts for disease control. *Wolbachia* from *Drosophila* and other mosquito species have been transferred to *Aedes aegypti* (which do not harbor *Wolbachia* naturally) and are now being released for arbovirus control around the world (Ross et al., 2019b). Several strains of *Wolbachia* block the transmission of arboviruses including dengue (Walker et al., 2011, Bian et al., 2010) and can induce cytoplasmic incompatibility, which reduces the viability of eggs produced by females which do not carry *Wolbachia* when they mate with males that do (Xi et al., 2005, Walker et al., 2011). Field trials deploying mosquitoes with the *w*Mel *Wolbachia* strain in Australia have established *Wolbachia*-carrying mosquitoes at stable high frequencies in the population (Hoffmann et al., 2011) and subsequently nearly eliminating local dengue transmission (O’Neill et al., 2018, Ryan et al., 2019). Later releases in dengue-endemic regions involving the *w*Mel or *w*AlbB strains of *Wolbachia* have reduced dengue cases by over 60% in some trial zones in Indonesia (Utarini et al., 2021), Malaysia (Hoffmann et al., 2024), Colombia (Velez et al., 2023) and Brazil (Pinto et al., 2021).

Population replacement programs of this nature where *Wolbachia* are introduced into populations involves the spread and persistence of *Wolbachia* in wild *Ae. aegypti* populations, which in turn depends on the fidelity of maternal transmission, strength of cytoplasmic incompatibility and host fitness costs induced by *Wolbachia* (Hoffmann and Turelli, 1997). Stable establishment of *Wolbachia* has been challenging in some environments (Hien et al., 2021, Gesto et al., 2021), which is likely driven by a complex set of factors, including characteristics of the released mosquito strain like pesticide resistance (Garcia et al., 2019), environmental effects on cytoplasmic incompatibility and maternal transmission (Ross et al., 2017b), as well as aspects of the built environment including dispersal barriers and heterogeneity in larval habitats (Schmidt et al., 2018, Hancock et al., 2019). Understanding which factors contribute to *Wolbachia* establishment could help guide decisions about where and when to release, the choice of *Wolbachia* strain, mass rearing procedures, and which locations may require supplementary releases or boosted numbers.

There is now a substantial body of evidence for host fitness costs of *Wolbachia* strains across the *Ae. aegypti* life cycle from egg to adult (Reviewed in Ross et al. (2019b), Figure 1). Many of these costs are both life stage-specific and context-dependent, occurring under conditions of environmental stress and amplified during quiescence and senescence. Costs strongly depend on *Wolbachia* strain (Walker et al., 2011, Fraser et al., 2017) and are influenced by genetic background (Carvalho et al., 2020, Ross and Hoffmann, 2022). There may also be interactions between the two factors, with studies in different mosquito backgrounds identifying contrasting patterns of fitness costs with the same strains (Joubert et al., 2016, Maciel-de-Freitas et al., 2024). On the other hand, many traits do not appear to be influenced by *Wolbachia* including mating success (Turley et al., 2013), host-seeking (Lau et al., 2020) and insecticide resistance (Endersby and Hoffmann, 2013).

**Figure 1.**
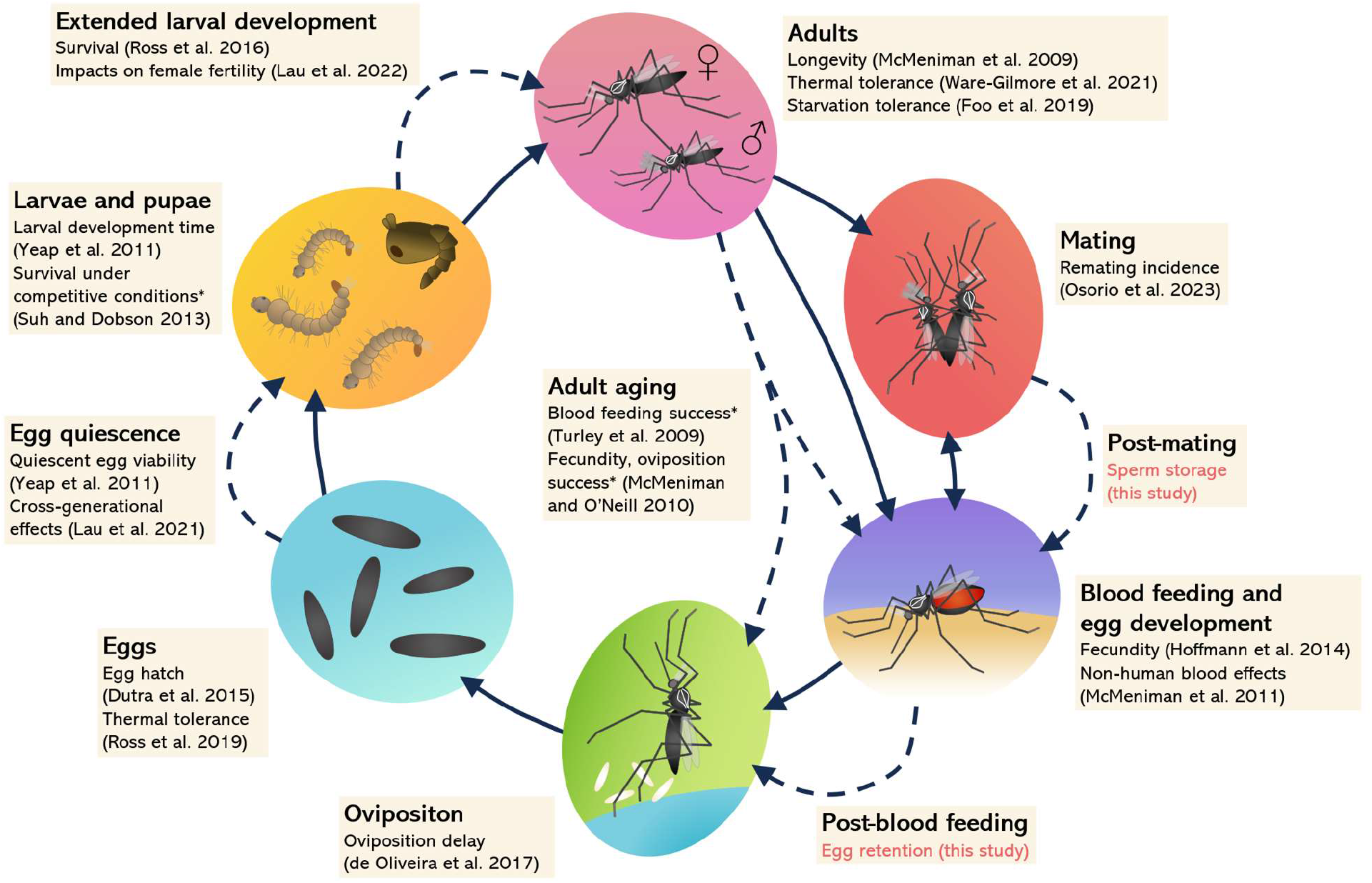
The life cycle of *Aedes aegypti* mosquitoes with examples of host fitness costs induced by *Wolbachia* strains for a range of traits. Solid lines indicate a progression across life stages, while dashed lines indicate an extension or pause at a particular stage such as adult aging and egg quiescence. Traits with * are only affected by the highly virulent *w*MelPop infection which is not currently being released in field trials. Note that costs are strain-dependent and may not be identified consistently across studies. Traits that have been tested and found to not be influenced by *Wolbachia* strains such as mating success are not included here except when tested in this study.

Because *Wolbachia* releases are occurring in a range of climates, some of them with limited rainfall, understanding the impacts of *Wolbachia* strains under a broad range of conditions including egg retention could be helpful for predicting its spread in the field. *w*AlbB and *w*MelM are two *Wolbachia* strains now being released for dengue control in different countries, including arid environments such as Jeddah, Saudi Arabia (Endersby-Harshman et al., 2021). *w*MelM was originally transferred from *Drosophila* melanogaster and is a variant of the widely released *w*Mel strain which is relatively more heat-resistant and has strong dengue blocking potential (Gu et al., 2022, Ross et al., 2023). Both the *w*AlbB and *w*MelM strains have impacts on fertility, particularly in quiescent states, and stress tolerance (Gu et al., 2022, Ross et al., 2023). We therefore measured the impact of both strains on the quantity and quality of eggs when females were forced to retain them under laboratory conditions. Furthermore, we investigated potential effects of *w*MelM on the quality of stored sperm given that females mating with males carrying *w*MelM have an increased remating frequency (Osorio et al., 2023), and that other *Wolbachia* strains in *Drosophila simulans* reduce sperm competitive ability (Champion de Crespigny and Wedell, 2006) and the ability of females to store sperm (Ferguson et al., unpublished). We find substantial impacts of *w*MelM but not *w*AlbB on egg hatch following extended egg retention. This effect of *w*MelM occurred regardless of whether females mated with *w*MelM or uninfected males, and some impacts persisted even when females laid a second batch of eggs without retention. In contrast, *w*MelM had no impact on long-term sperm storage, with declines in fertility largely driven by female age. While the importance of egg retention in the wild has yet to be quantified, the impact of *Wolbachia* on this trait may influence the spread of *Wolbachia* and help to explain heterogeneity in invasion success in some environments.

## Methods

### Ethics statement

Blood feeding of mosquitoes on human volunteers was approved by the University of Melbourne Human Ethics committee (approval 0723847). All adult subjects provided informed written consent (no children were involved).

### Mosquito populations and rearing

We used three *Aedes aegypti* populations in this study on a common (North Queensland, Australia) genetic background. Mosquitoes carrying the *w*MelM variant of *Wolbachia* were generated through microinjection of cytoplasm from field-collected *D. melanogaster* as described previously (Gu et al., 2022). Mosquitoes carrying the *w*AlbB strain (*w*AlbB-Hou variant) were generated through microinjection of the strain generated by Xi et al. (2005) to *Aedes aegypti* with an Australian genetic background (Ross et al., 2021). Uninfected mosquitoes were generated by curing the *w*MelM population of *Wolbachia* by treating adults with 2 mg mL^−1^ tetracycline hydrochloride in a 10% sucrose solution across two consecutive generations. Females from all three populations were backcrossed to males from a naturally uninfected laboratory population originating from Cairns, North Queensland, for two consecutive generations prior to the experiments. Mosquitoes for colonies and all experiments were maintained under controlled laboratory conditions at 26°C with a 12:12 light:dark cycle according to methods described previously (Ross et al., 2017a).

To rear mosquitoes for experiments, eggs from colonies (< 2 weeks old) were hatched in trays with reverse osmosis (RO) water and a few grains of yeast. First instar larvae were transferred to trays with 4 L of RO water at a density of 400 larvae per tray and provided with Hikari tropical sinking wafers (Kyorin food, Himeji, Japan) *ad libitum*. Pupae were sexed and allowed to emerge into separate cages (19.7-L BugDorm-1, MegaView Science Co., Ltd., Taichung City, Taiwan) before establishing crosses (see below) when adults were 2-3 d old. Adults were provided with water and a 10% sucrose solution until 1 d before blood feeding, when sugar was removed.

### Egg retention and oviposition

Methods for forcing egg retention were adapted from a previous study (Venkataraman et al., 2023). Female mosquitoes (5-6 d old) were blood fed on the forearm of a human volunteer (University of Melbourne human ethics approval 0723847) and transferred to BugDorm-1 cages provided with only sugar through a cotton wick (Figure 2). Oviposition sites were removed and any water droplets within the cage were wiped up. Open containers of water were placed on top of each cage and cages were then placed within plastic bags. This procedure largely prevented mosquitoes from laying eggs while maintaining a high humidity to reduce mortality. For each treatment, we set up at least three cages (with approximately 200 females and 200 males each) and only used mosquitoes from cages where no eggs were visible. In some cases, mosquitoes laid a small number of eggs on the sugar wick and these cages were discarded.

**Figure 2.**
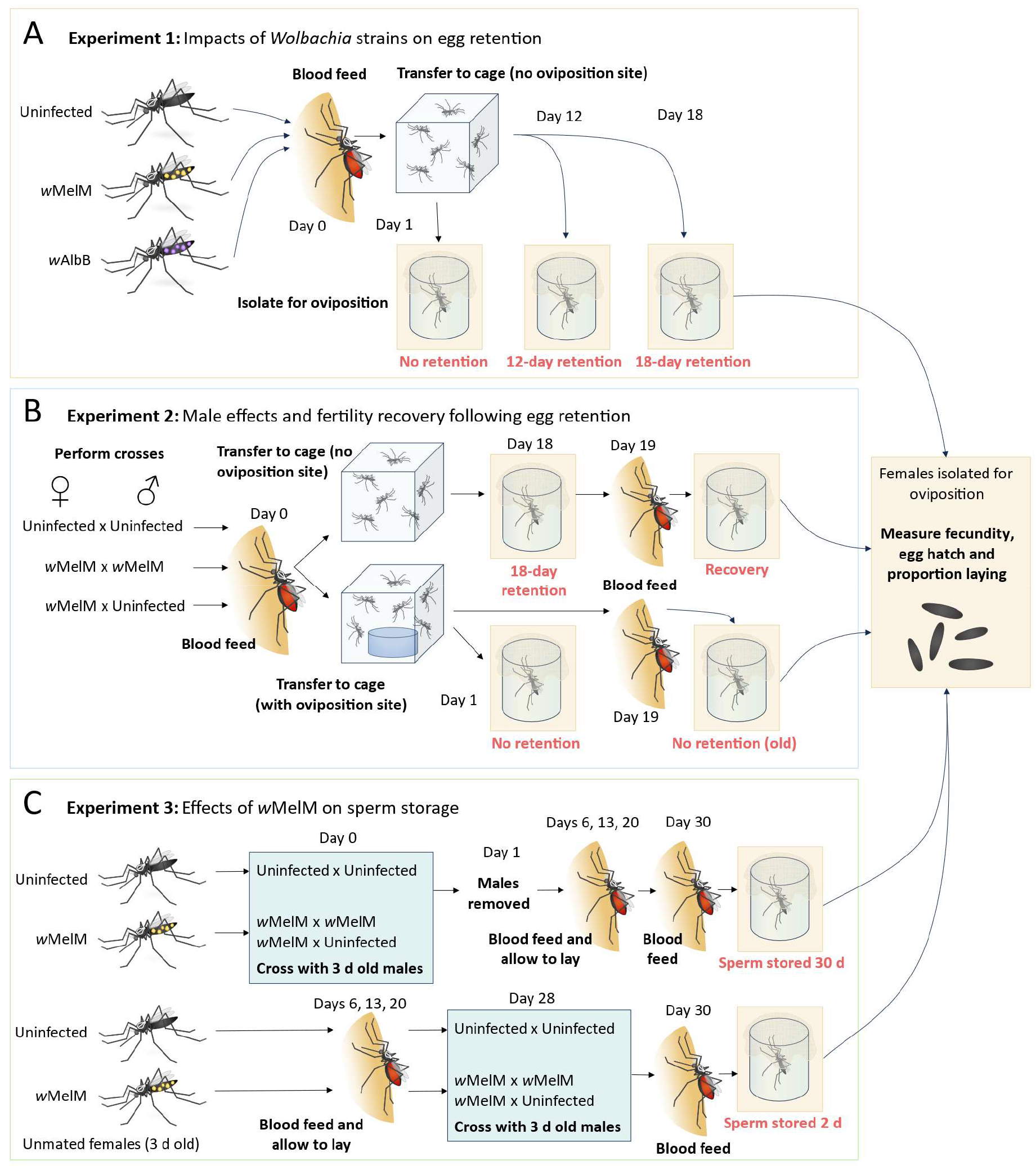
Experimental design. In Experiment 1 (A), mated females from the uninfected, wMelM and wAlbB populations experienced no egg retention or 12 or 18 days of egg retention before laying eggs. In Experiment 2 (B), *w*MelM and uninfected females were crossed in different combinations. Females then either experienced no egg retention or 18 days of egg retention. Females from both groups were then blood fed again to initiate a second gonotrophic cycle. In Experiment 3 (C), uninfected and *w*MelM females were crossed to 3 d old males when they were either 3 or 31 d old, then blood fed and isolated for oviposition either 30 or 2 d after mating. Text in red indicates the treatments measured in each experiment for each *Wolbachia* strain or cross. All treatments were measured for fecundity, egg hatch proportion and the proportion of females laying eggs. For experiments 1 and 2, Day 0 corresponds to the day of first blood feeding, while for Experiment 3 Day 0 corresponds to when mosquitoes were first mated.

Female mosquitoes from egg retention cages were isolated in 70 mL specimen cups at different time points post-blood feeding (Figure 2). Cups were filled with 20 mL of larval rearing water, lined with strips of sandpaper (Norton Master Painters P80 sandpaper, Saint-Gobain Abrasives Pty. Ltd., Thomastown, Victoria, Australia), and covered with a mesh lid to prevent mosquitoes from escaping. Cups were checked daily for 3 days and sandpaper strips were collected from females that laid eggs. We recorded the number of females that died as well as the number that did not lay eggs after 3 days. Sandpaper strips were partially dried on paper towel then placed in sealed containers with a layer of paper towel for 3 d. Eggs were hatched by filling trays with water and adding a few grains of yeast. The next day, the number of unhatched and hatched eggs (with a clearly detached cap) were counted to determine fecundity and egg hatch for individual females.

### Experiment 1 – Impacts of Wolbachia strains on egg retention

In the first experiment (Figure 2A), we measured the impact of egg retention on fecundity and egg hatch proportions for the uninfected, *w*MelM and *w*AlbB populations. Males and females were crossed in groups in cages (i.e. mass mated with approximately 200 females and 200 males per cage) within populations only. Females were kept in cages with no access to oviposition sites for 1 (no retention), 12 or 18 d post-blood feeding before isolating them for oviposition. We set up thirty replicate cups for the 1 and 12 d time points and 20 replicates for the 18 d time point.

### Experiment 2 – Male effects and fertility recovery following egg retention

In the second experiment (Figure 2B), we compared the uninfected and *w*MelM populations again and included an additional cross and treatment. Our aims were to: 1. Test whether the costs of *w*MelM to egg retention were driven by females or males; and 2. Test the extent of fertility recovery in a subsequent gonotrophic cycle after extended egg retention. Uninfected mosquitoes were crossed in groups within the line, and *w*MelM females were crossed to either *w*MelM males or uninfected males in groups. After blood feeding, half the mosquitoes were transferred to egg retention cages with no oviposition site then isolated for egg laying at 18 d post-blood feeding. To test for recovery, females that survived after 18 d of egg retention (24-25 d old) were blood fed again to collect their eggs from a second gonotrophic cycle. The other half of the mosquitoes were transferred to colony cages containing an oviposition site. Females from this cage were isolated for oviposition 1 d later as a “no egg retention control”, and the remainder were allowed to lay eggs freely. These mosquitoes were blood fed again at 24-25 d old and isolated to collect eggs from a second gonotrophic cycle. We set up 60 replicates per cross for the two controls (5-6 d old females with no egg retention, and 24-25 d old females with no egg retention in either gonotrophic cycle), and 120 replicates for the 18 d egg retention crosses. All females that survived the 18 d egg retention treatment were blood fed and isolated again for a second gonotrophic cycle, though initial sample sizes were lower than 120 due to mortality. In this experiment, we recorded the proportion of surviving females that laid eggs-those that died were excluded from the analysis.

### Experiment 3 – Effects of wMelM on sperm storage

In the third experiment (Figure 2C), we compared the fertility of females that were either mated when they were 3 days old or 31 days old, then blood fed at 33 days old. Females in all treatments were blood fed four times, with females isolated following the fourth blood meal to measure fertility. In the sperm storage treatment (sperm stored 30 d), females were given 24 hrs to mate with males in groups then all males were removed to ensure that they could not remate for the next 30 d. In the second treatment (sperm stored 2 d), unmated females were kept separate from males and then mated to 3 d old males in groups for 24 hrs. In both treatments, uninfected mosquitoes were crossed within the line and *w*MelM females were crossed to either *w*MelM males or uninfected males. Effects of long-term sperm storage were measured by comparing the fecundity and egg hatch of females that were blood fed either 2 d or 30 d after mating with males.

### Statistics

All statistical analyses were performed in SPSS Statistics version 29.0.0.0. For all experiments, we used general linear models to test for effects of *Wolbachia* strain, cross and egg retention or sperm storage treatment and their interactions on fecundity and egg hatch proportions (which were logit transformed for normality (Warton and Hui, 2011)). Females that died or laid no eggs were excluded from the analyses of fecundity and egg hatch. Tukey’s post-hoc with multiple comparisons were used to compare treatments within a cross or *Wolbachia* strain. For experiments 2 and 3, we performed an initial analysis with only the *w*MelM female x *w*MelM male and *w*MelM female x uninfected male crosses to test whether male strain had any influence on fecundity and egg hatch following egg retention or sperm storage. Given the absence of male effects, we then pooled these two crosses for additional analyses focused on testing the effects of *w*MelM in the female. In Experiment 2, where large sample sizes were available, we also computed the proportions of mosquitoes laying eggs in each treatment, which we analysed with Fisher’s exact tests treating each female as a data point. We calculated the total number of viable offspring per female by multiplying the number of eggs laid by their hatch proportion. By also including data from females that failed to lay any eggs, this measure provides an overall indication of the total fitness effects of the infection.

## Results

### Impacts of Wolbachia strain on egg retention

To test for effects of *Wolbachia* strain on egg retention, eggs from uninfected, *w*MelM and *w*AlbB populations were collected and hatched after females experienced 0, 12 or 18 d of forced egg retention (Figure 2A). We found significant effects of egg retention treatment on both fecundity (GLM: F_2,208_ = 17.309, P < 0.001) and egg hatch proportion (F_2,208_ = 38.654, P < 0.001), where both traits in all populations experienced a decline with extended egg retention (Figure 3). For fecundity, there was no significant effect of *Wolbachia* strain (F_2,208_ = 1.657, P = 0.193) or a significant interaction between *Wolbachia* strain and egg retention (F_4,208_ = 1.019, P = 0.399). In contrast, egg hatch proportions were influenced by *Wolbachia* strain (F_2,208_ = 40.170, P < 0.001) and there was an interaction between *Wolbachia* strain and egg retention treatment (F_4,208_ = 5.747, P = 0.002).

**Figure 3.**
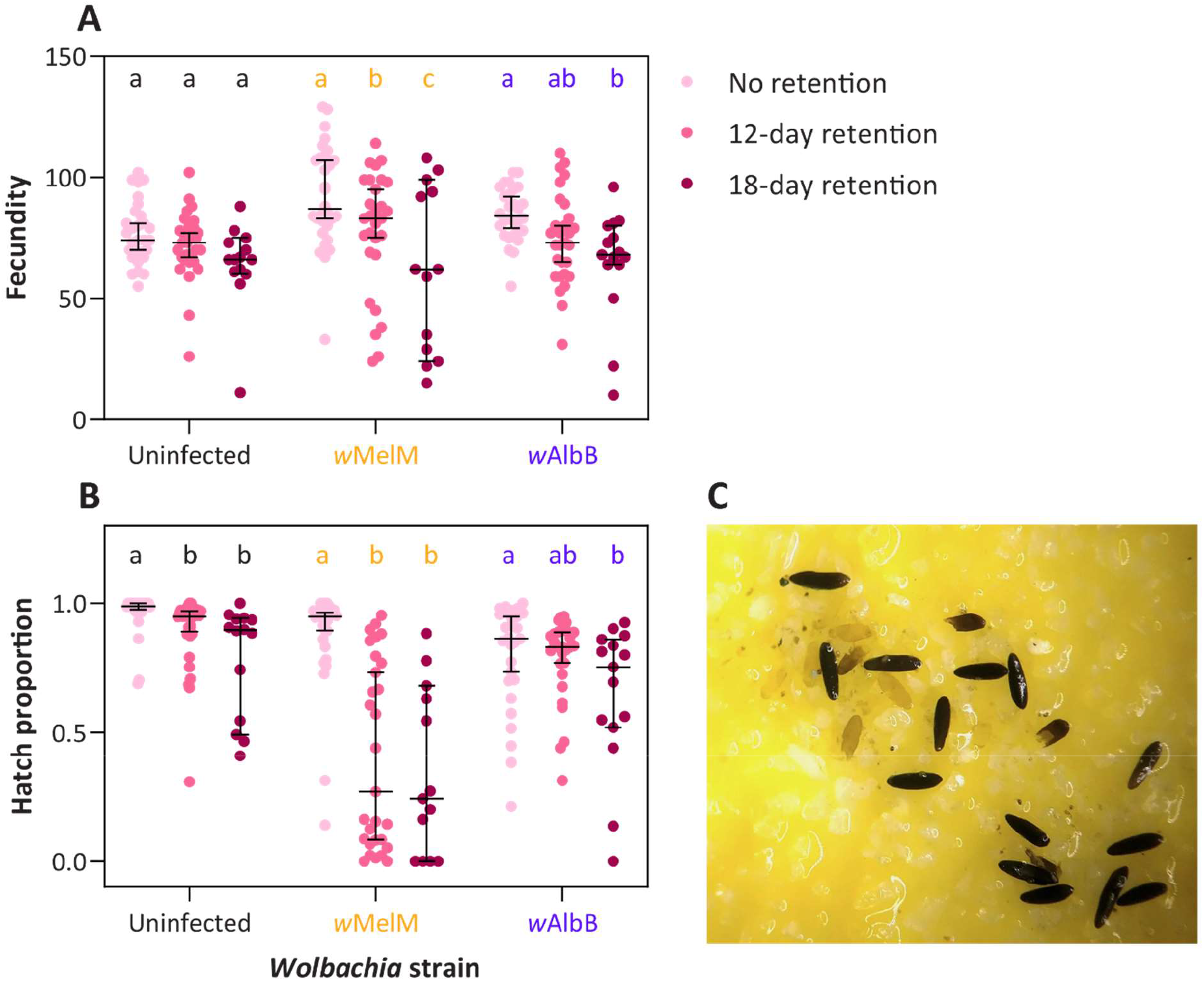
Costs of *Wolbachia* to the quality of retained eggs depend on *Wolbachia* strain. (A) Fecundity and (B) hatch proportions of eggs laid by uninfected, *w*MelM or *w*AlbB populations of *Aedes aegypti* after 0, 12 or 18 days of forced retention. Dots show data for individual females while horizontal lines and error bars show medians and 95% confidence intervals. Within each population, different letters represent significant differences (P < 0.05) between egg retention treatments based on Tukey’s post-hoc tests with a correction for multiple comparisons. (C) Example of egg defects observed in eggs laid by *w*MelM females after extended egg retention. Note the mis-formed and unmelanized eggs laid on the oviposition substrate.

The *w*MelM females experienced a substantial decline in egg hatch proportion following egg retention, with a median egg hatch proportion that was 71.5% lower than the control after 12 d and 74.5% lower after 18 d (Figure 3). While *w*AlbB females had lower egg hatch proportion overall compared to uninfected females, the percent decline in hatch proportion with egg retention was similar (9.4% for uninfected and 12.8% for *w*AlbB after 18 d). These results suggest a strain-specific effect of *Wolbachia* on the quality of retained eggs, with greatly reduced egg hatch proportion for *w*MelM but not *w*AlbB. During this experiment we noticed that many *w*MelM females (e.g. 11/20 at 18 d) produced no or few viable offspring, and inspection under a dissecting microscope revealed defects with eggs that were misshapen or unmelanized (Figure 3C provides an example).

### Male effects on egg retention and fertility recovery

We were interested in testing if costs of *w*MelM to retained egg quality may be mediated by infections in both the male and female and if these could potentially recover in subsequent gonotrophic cycles. We therefore tested the impact of *w*MelM on retained egg quality when females were crossed to either *w*MelM or uninfected males. We also tested whether fertility recovered following an additional gonotrophic cycle without egg retention. We first analysed the data from *w*MelM females that were crossed to *w*MelM or uninfected males. While there was a substantial effect of egg retention treatment on fecundity (GLM: F_3,353_ = 31.410, P < 0.001) and egg hatch proportion (F_3,353_ = 148.878, P < 0.001), the male strain used in crosses had no significant effect on either trait (fecundity: F_1,353_ = 0.1.026, P = 0.749, egg hatch: F_1,353_ = 2.924, P = 0.088, Figure S1). However, there was a significant interaction between male strain and egg retention treatment for egg hatch (F_3,353_ = 4.921, P = 0.002) though not for fecundity (F_3,353_ = 0.179, P = 0.911).

We then pooled data from different male strains to analyze effects of *Wolbachia* strain on egg retention in females. We found significant effects of female strain (GLM: F_1,620_ = 304.010, P < 0.001), egg retention treatment (F_3,620_ = 42.310, P < 0.001) and an interaction between strain and treatment (F_3,620_ = 8.027, P < 0.001) for fecundity. The fecundity of females laying retained eggs declined for *w*MelM but not for uninfected females compared to females in the control with no egg retention (Figure 4A). When comparing the two treatments where females were 24-25 d old and in their second gonotrophic cycle, uninfected females laid more eggs when they previously experienced egg retention, while *w*MelM females laid fewer eggs (Figure 4A).

**Figure 4.**
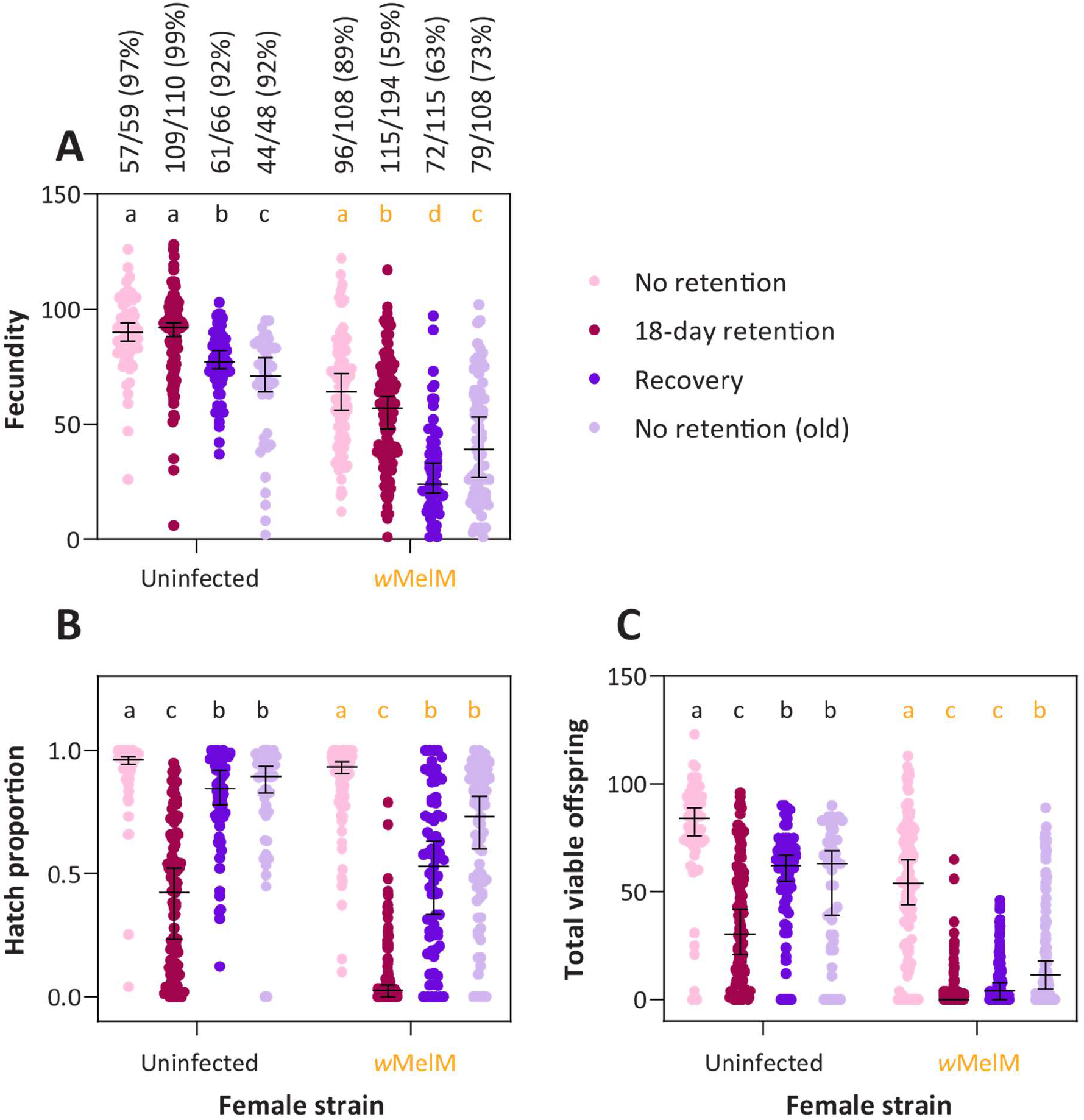
Fertility of *w*MelM females only partially recovers from extended egg retention. (A) fecundity, (B) hatch proportions and (C) total viable offspring from uninfected and *w*MelM females. Females were blood fed at 5-6 d old and experienced no retention (pink) or 18 days of egg retention (maroon). Females were also blood fed at 24-25 d old following 18 d of egg retention (dark purple) or no egg retention (light purple) for an additional gonotrophic cycle. For the full experimental design see Figure 2B. Data for *w*MelM are pooled from crosses with both uninfected and *w*MelM males (see Figure S1 for male data). Numbers and percentages in panel A indicate the number and percentage of females in each treatment that laid eggs out of the total number surviving. Dots show data for individual females while horizontal lines and error bars show medians and 95% confidence intervals. <u>Within</u> each strain, different letters represent significant differences (P < 0.05) between egg retention treatments according to Tukey’s post-hoc tests with a correction for multiple comparisons.

The proportion of females laying eggs did not differ significantly between *w*MelM and uninfected females in the control (Fisher’s exact test: P = 0.142), but for females in the egg retention treatment significantly fewer *w*MelM females laid eggs (P < 0.001, Figure 4A). There was also an apparent effect of age, with decreased proportions of 24-25 d old *w*MelM females laying eggs, even for females that experienced no retention (P = 0.010, Figure 4A). There was minimal mortality of isolated females across the experiment, except for the recovery treatment where only 66/110 (60%) of uninfected and 115/194 (59.3%) of *w*MelM females survived after a second blood meal.

We also found significant effects of strain (GLM: F_1,620_ = 84.880, P < 0.001), treatment (F_3,620_ = 182.007, P < 0.001) and an interaction (F_3,620_ = 11.073, P < 0.001) on (logit transformed) egg hatch proportions (Figure 4B). Egg hatch proportion declined in both strains following extended egg retention, but to a much greater extent for *w*MelM females (median 0.026) than for uninfected females (median 0.422) (Figure 4B). Egg hatch proportion recovered in the following gonotrophic cycle for both strains, resulting in proportions that were similar to those obtained with females of the same age that experienced no egg retention (Figure 4B).

Egg retention substantially reduced total offspring counts (which included females that laid no eggs), particularly for *w*MelM females where median offspring declined to zero (Figure 4C). The decline in offspring count with female age was also particularly pronounced for *w*MelM females (Figure 4C). However, effects of *w*MelM were not due to age alone since egg hatch proportions, total offspring counts and the proportion of females laying eggs was lower in the egg retention treatment compared to old females that experienced no egg retention (Figure 4). These patterns indicate a substantial impact of *w*MelM on fertility with both age and egg retention, with sustained impacts even when completing a second gonotrophic cycle without retention.

### Effects of wMelM on sperm storage

In the previous experiment, *w*MelM had negative effects on fertility particularly in older females. While this may reflect a deterioration of female fertility with age, costs may also be associated with sperm quality given that females use stored sperm in subsequent gonotrophic cycles. We therefore tested for impacts of *w*MelM on sperm storage by comparing the fecundity and egg hatch of females that were blood fed either 2 d or 30 d after mating. Crosses involving both *w*MelM and uninfected mosquitoes allowed us to separate effects of *w*MelM in males on stored sperm quality and the effect of *w*MelM in females on sperm storage.

When considering only *w*MelM females that mated with either *w*MelM or uninfected males, we found an effect of male strain (GLM: F_1,72_ = 8.976, P = 0.004), sperm storage treatment (F_1,72_ = 7.149, P = 0.009) and an interaction (F_1,72_ = 4.843, P = 0.031) between these two factors on fecundity (Figure S2). Fecundity was higher in females using sperm stored for 2 d compared to females using sperm stored for 30 d at the point of blood feeding, with this difference being more pronounced when females were crossed to uninfected males (Figure S2). In contrast, there was no significant effect of any factor or interaction on egg hatch proportion (all P > 0.095, Figure S2).

When comparing the effects of female strain with data pooled across *w*MelM and uninfected males, we found that *w*MelM significantly reduced fecundity (F_1,118_ = 70.42, P < 0.001) and (logit transformed) egg hatch proportion (F_1,118_ = 19.15, P < 0.001, Figure 5). Females with sperm stored for 30 d had a lower fecundity compared to females with sperm stored for 2 d (F_1,118_ = 13.94, P < 0.001) but the hatch proportions of their eggs were higher (F_1,118_ = 15.45, P < 0.001). Patterns with respect to sperm storage treatment were consistent across both wMelM and uninfected females (Figure 5) with no significant interaction between sperm storage treatment and female strain for either fecundity (F_1,118_ = 0.092, P = 0.762) or egg hatch proportion (F_1,118_ = 3.065, P = 0.083), suggesting that *w*MelM has no deleterious effect on the quality of stored sperm specifically.

**Figure 5.**
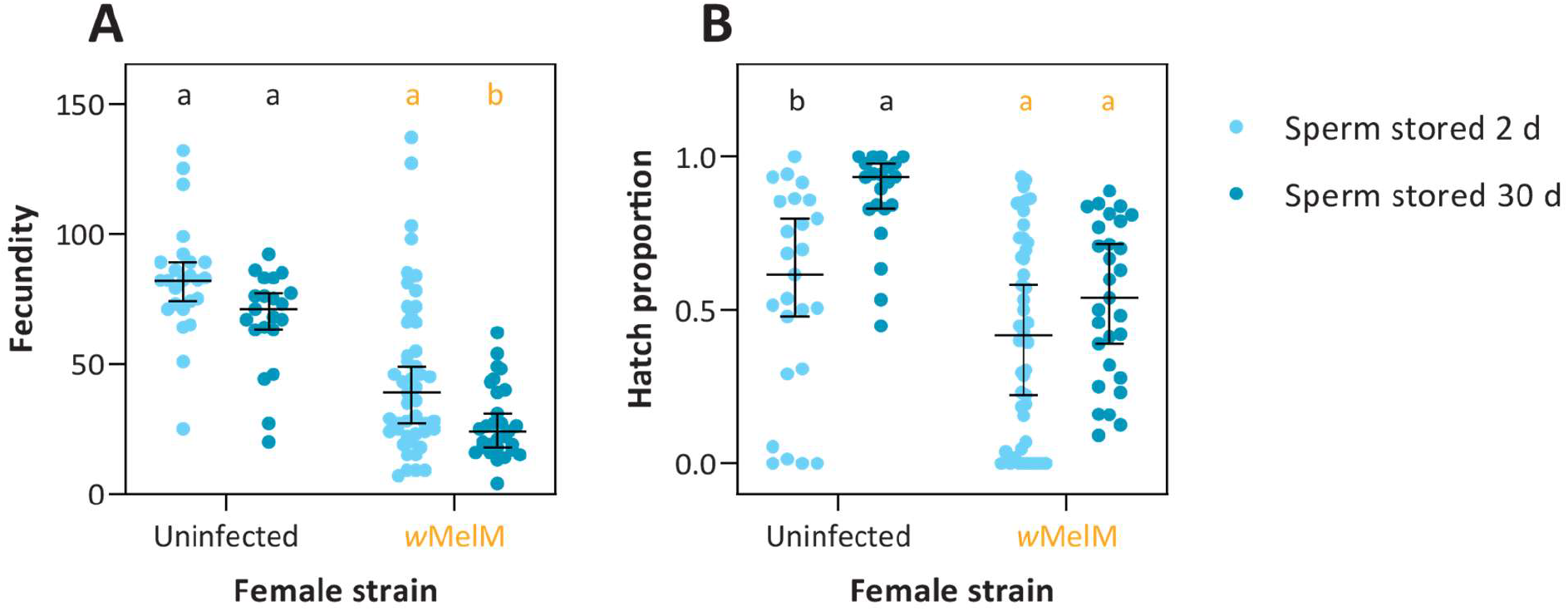
Costs of *w*MelM to fecundity and egg hatch are unaffected by sperm storage. (A) Fecundity and (B) hatch proportions of eggs laid by uninfected or *w*MelM females crossed to 3 d old males when they were either 3 or 31 d old, then blood fed 2 d (light blue) or 30 d (dark blue) after mating respectively. For the full experimental design see Figure 2C. Dots show data for individual females while horizontal lines and error bars show medians and 95% confidence intervals. Within each strain, different letters represent significant differences (P < 0.05) between sperm storage treatments according to GLMs with Tukey’s post-hoc tests involving a correction for multiple comparisons.

## Discussion

We show that *Wolbachia* strain *w*MelM disrupts the ability of *Ae. aegypti* females to retain viable eggs for extended periods, with a decrease in the proportion of females that lay, as well as a substantial reduction in the proportion of those eggs that hatch. The phenotype described here adds to a growing list of traits influenced by *Wolbachia* strains in *Ae. aegypti* (Figure 1), and is consistent with other studies that have described context-dependent and strain-dependent fitness costs, particularly to traits related to fertility (Lau et al., 2021). The costs described here and in other studies could contribute to challenges in establishing this strain in field populations, especially in climates with intermittent rainfall where mosquitoes may frequently retain eggs.

Our comparison of the *w*AlbB and *w*MelM strains shows that costs of *Wolbachia* are strain-specific, since no clear effects of *w*AlbB on fecundity or egg hatch following retention were identified. The lack of effect of *w*AlbB was surprising given that substantial costs of this strain have been identified previously, particularly when mosquito eggs are in quiescent states (e.g. (Axford et al., 2016, Lau et al., 2021), though other studies do show higher costs of a different *w*Mel variant compared to wAlbB under some conditions (Joubert et al., 2016, Maciel-de-Freitas et al., 2024). Retained eggs laid by *w*MelM females often had low hatch proportions, which may be partially explained by the presence of defects such as incomplete melanization, which were rarely detected in the uninfected and *w*AlbB populations. These defects are similar to those of eggs laid by females carrying the *w*MelPop *Wolbachia* strain when females are aged or fed on non-human blood (McMeniman et al., 2011), but whether a common mechanism is involved remains to be explored.

Under some conditions, such as nutrient deprivation, *Ae. aegypti* females can resorb their oocytes (Lea et al., 1978, Clifton and Noriega, 2012). In our experiments, we found a minimal decline in fecundity for uninfected females following extended egg retention. In contrast, *w*MelM females tended to lay fewer eggs under the same conditions and many females laid no eggs. This may reflect females resorbing their eggs or possibly continued retention of eggs despite having access to an oviposition site, which could be explored through dissections. Our experiments show that costs of *w*MelM were driven mainly by the female since effects occurred regardless of the male used in the crosses. Our sperm storage experiments also suggest that *w*MelM has no impact on sperm quality/sperm viability, or on female ability to store sperm, indicating that costs of *w*Mel relate to the quality of retained eggs or the ability of females to fertilize eggs, which occurs during oviposition (Degner and Harrington, 2016a).

We acknowledge that some of the effects of *w*MelM described in our study could be due to mosquito age rather than solely due to egg retention effects. Previous studies have identified costs of *Wolbachia* to fertility that increase with age (McMeniman and O’Neill, 2010), and we show costs here for *w*MelM when females of different ages (but with no egg retention) are compared. However, total offspring counts for *w*MelM females were lower in the retention treatment compared to those for older females that experienced no retention, highlighting that effects extend beyond age. The effects of *w*MelM on egg retention do also appear to persist to some extent since total offspring counts did not fully recover after feeding again, in contrast to uninfected mosquitoes which fully recovered.

In summary, our study describes substantial costs of *w*MelM, a *Wolbachia* strain now being released in dengue control programs, to a trait which is likely to be important in environments with variable rainfall. While the experimental conditions used here might be regarded as relatively extreme, the fact that *Aedes aegypti* are capable of extended egg retention while retaining viability does suggest that it may link to an adaptive physiological trait. Furthermore, while costs of *Wolbachia* to individual traits can be minor, the cumulative costs across many traits (Figure 1) can add up to be quite substantial under some conditions (Ross et al., 2023), which could explain fluctuations and loss of *Wolbachia* in some field release sites (Hien et al., 2021, Nazni et al., 2019). While we did not consider other environmental factors such as temperature, these may also interact with effects on egg retention leading to additional costs. The costs described here and elsewhere also raise important evolutionary questions, since successful *Wolbachia* establishment will result in mosquito populations that are less tolerant of dry environments if their ability to retain eggs is disrupted. If *Wolbachia* remains at high frequency these costs may shift over time as recently demonstrated for quiescent egg viability (Ross and Hoffmann, 2022). These environment-specific effects raise issues about the long-term success of the widespread replacement of wild populations with mosquitoes carrying *Wolbachia* strains for dengue reduction under climate change (Ross and Hoffmann, 2023). It may be that the suitability of different strains changes under more variable future rainfall patterns particularly if temperature extremes lead to higher evapotranspiration rates.

## Supplementary information

**Figure S1.**
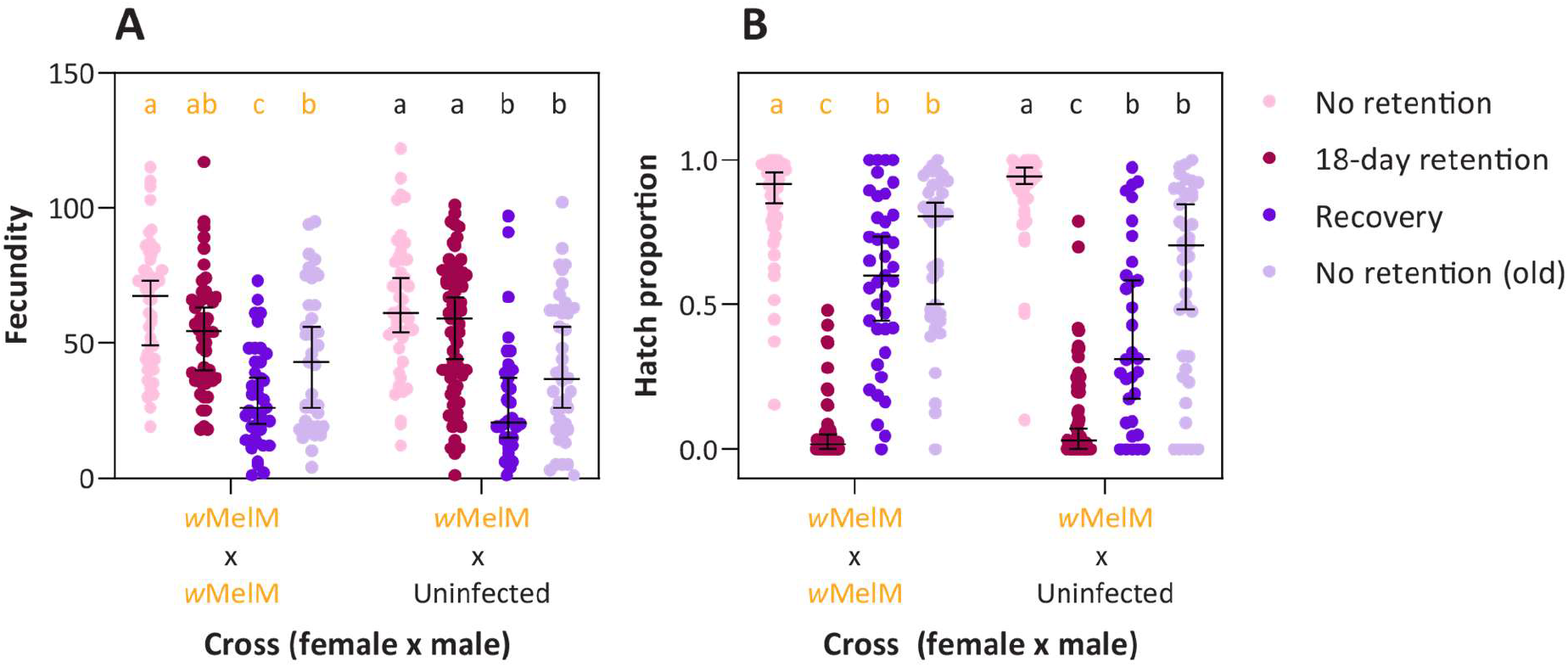
Male effects of *w*MelM on the quality of retained eggs. (A) Fecundity and (B) hatch proportions of eggs laid by *w*MelM females after mating with *w*MelM or uninfected males. Females were blood fed at 5-6 d old and experienced no retention (pink) or 18 days of egg retention (maroon). Females were also blood fed at 24-25 d old following 18 d of egg retention (dark purple) or no egg retention (light purple) for an additional gonotrophic cycle. For the full experimental design see Figure 2B. Dots show data for individual females while horizontal lines and error bars show medians and 95% confidence intervals. Within each cross, different letters represent significant differences (P < 0.05) between egg retention treatments based on Tukey’s post-hoc tests with a correction for multiple comparisons.

**Figure S2.**
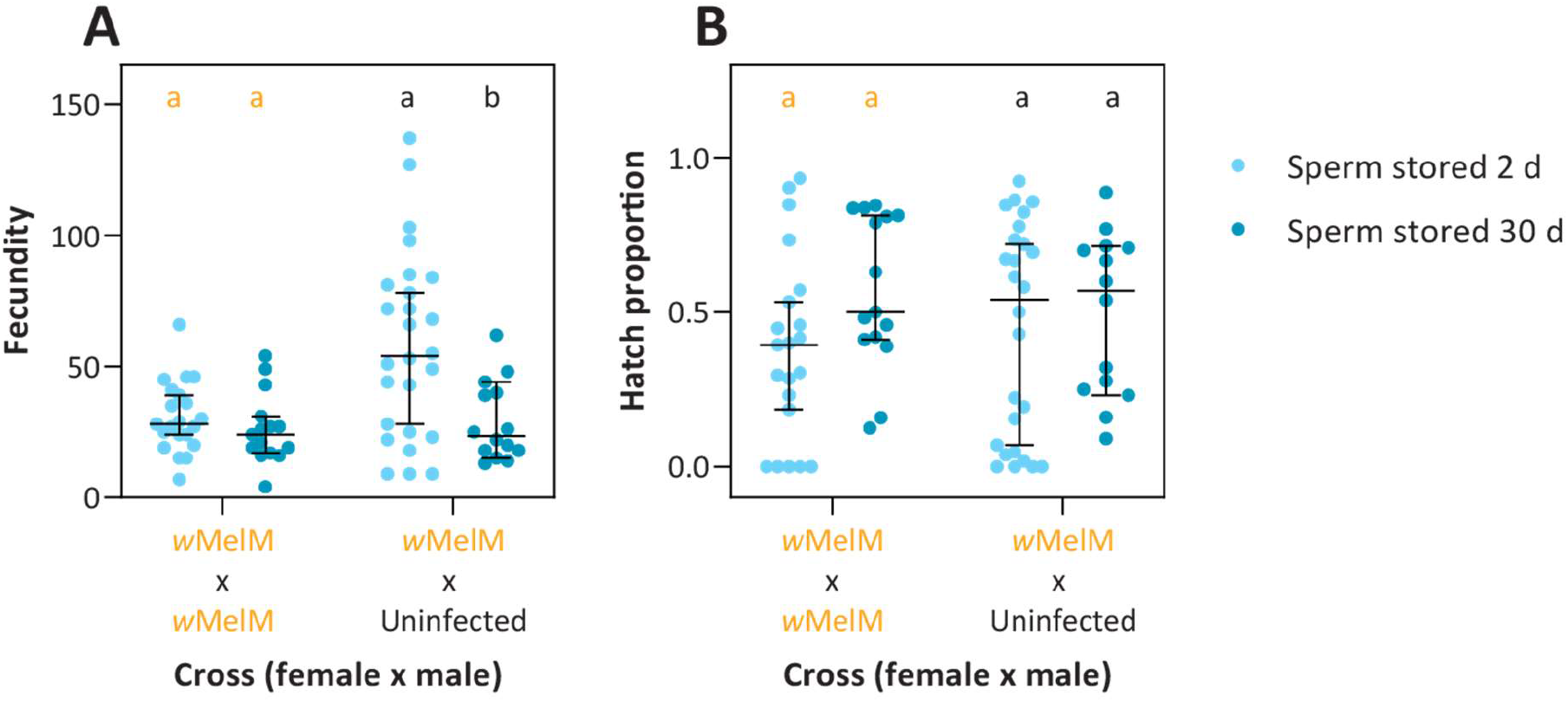
Male effects of wMelM on fecundity and egg hatch following sperm storage. (A) Fecundity and (B) hatch proportions of eggs laid by *w*MelM females after mating with *w*MelM or uninfected males. *w*MelM females were crossed to 3 d old males when they were either 3 or 31 d old, then blood fed 2 d (light blue) or 30 (dark blue) after mating respectively. For the full experimental design see Figure 2C. Dots show data for individual females while horizontal lines and error bars show medians and 95% confidence intervals. Within each cross, different letters represent significant differences (P < 0.05) between sperm storage treatments based on Tukey’s post-hoc tests with a correction for multiple comparisons.

